# mRNA Treatment Rescues Niemann-Pick Disease Type C1 in Patient Fibroblasts

**DOI:** 10.1101/2022.02.21.479058

**Authors:** Denzil Furtado, Christina Cortez-Jugo, Ya Hui Hung, Ashley I. Bush, Frank Caruso

## Abstract

Messenger RNA (mRNA) holds great potential as a disease-modifying treatment for a wide array of monogenic disorders. Niemann-Pick disease type C1 (NP-C1) is an ultra-rare monogenic disease that arises due to loss-of-function mutations in the *NPC1* gene, resulting in the entrapment of unesterified cholesterol in the lysosomes of affected cells and a subsequent reduction in their capacity for cholesterol esterification. This causes severe damage to various organs including the brain, liver, and spleen. In this work, we describe the use of NPC1-encoded mRNA to rescue the protein insufficiency and pathogenic phenotype caused by biallelic *NPC1* mutations in cultured fibroblasts derived from an NP-C1 patient. We first evaluated engineering strategies for the generation of potent mRNAs capable of eliciting high protein expression across multiple cell types. We observed that “GC3” codon optimization, coupled with N1-methylpseudouridine base modification, yielded an mRNA that was approximately a thousand-fold more potent than wildtype, unmodified mRNA in a luciferase reporter assay, and consistently superior to other mRNA variants. Our data suggest that the improved expression associated with this design strategy was due in large part to the increased secondary structure of the designed mRNAs. Both codon optimization and base modification appear to contribute to increased secondary structure. Applying these principles to the engineering of NPC1-encoded mRNA, we observed a normalization in NPC1 protein levels after mRNA treatment, as well as a rescue of the mutant phenotype. Specifically, mRNA treatment restored the cholesterol esterification capacity of patient cells to wildtype levels, and induced a significant reduction in both unesterified cholesterol levels (>57% reduction compared to Lipofectamine-treated control in a cholesterol esterification assay) and lysosome size (157 μm^2^ reduction compared to lipofectamine-treated control). These findings show that engineered mRNA can correct the deficit caused by *NPC1* mutations. More broadly, they also serve to further validate the potential of this technology to correct diseases associated with loss-of-function mutations in genes coding for large, complex, intracellular proteins.

## Introduction

Niemann-Pick disease type C (NP-C) is a rare inherited neurovisceral disorder that arises due to inactivating mutations in one of two protein-coding genes, *NPC1* (95% of cases) or *NPC2*.^1^ The incidence of NP-C1 disease is approximately 1/90,000 live births, but due to the extreme heterogeneity of clinical phenotypes, a late-onset form of NP-C1 is estimated to have a much higher incidence rate of between 1/19,000–1/36,000.^2^ Both NPC1 and NPC2 are endo-lysosomal proteins that, under normal circumstances, act cooperatively to shuttle unesterified cholesterol obtained from endocytosed lipoproteins across the late endosomal membrane into the cytosol.^3^ NPC2, a small, soluble protein, binds unesterified cholesterol in the lumen and presents it to the N-terminal domain of NPC1.^4,5^ NPC1, which by contrast is a large, multipass transmembrane protein, subsequently facilitates cholesterol egress across the endosomal membrane.^4,6^

In NP-C disease, however, defective trafficking leads to the sequestration and accumulation of unesterified cholesterol and glycosphingolipids in lysosomes of cells throughout the body. This results in damage to various organs including the central nervous system (CNS), liver, spleen, and sometimes the lungs.^1^ The clinical manifestations of these pathologies often arise at different times and follow very different trajectories.^7^ In contrast, some severe infantile forms of the disease result in fatal liver or respiratory failure,^7^ while other adult forms result in late-onset neurodegenerative disease.^2^ Nevertheless, classical neurological manifestations in the majority of patients include vertical supranuclear gaze palsy, cerebellar ataxia, dystonia, dysarthria, and dysphagia, gelastic cataplexy, and epileptic seizures.^1,2,7^ The severity of these symptoms tends to worsen with time, since NP-C disease is chronic, progressive, and invariably fatal.^8^

Despite many and varied efforts, few treatments for NP-C disease exist.^9^ No drug has yet been approved by the United States (US) Food and Drug Administration (FDA), although various compounds have been introduced into late-stage clinical trials, including arimoclomol, an inducer of heat shock proteins (ClinicalTrials.gov identifier NCT02612129);^10^ miglustat, a glucosylceramide synthase inhibitor (ClinicalTrials.gov identifier NCT01760564);^11,12^ and hydroxypropyl-beta-cyclodextrin, a cyclic oligosaccharide that has been shown to sequester and transport unesterified cholesterol from the late endosome to the cytosol, thereby enabling increased cholesterol esterification (ClinicalTrials.gov identifier NCT04860960).^13^ Miglustat is currently approved in the EU, Canada and Japan for the treatment of progressive neurological complications in NP-C disease, and is frequently prescribed off-label for NP-C1 disease in the US.^12^ Hydroxypropyl-beta-cyclodextrin, meanwhile, has been approved under compassionate use protocols to treat several NP-C disease patients worldwide since 2009.^14^ Even with the aforementioned drugs, however, the prognosis for patients remains poor and the disease burden remains high.

Since NP-C disease occurs due to a lack of properly functioning NPC1 or NPC2 protein, therapeutic approaches that introduce a functional copy of the affected protein have been explored. Protein replacement therapy is not available for NP-C1 disease since it involves a transmembrane protein, but has demonstrated partial efficacy in a mouse model of NP-C2 disease.^15^ Viral gene therapy strategies, reliant on adeno-associated viral vector serotype 9 (AAV9), have shown promise in mouse models of NP-C1 disease when administered systemically^16^ and locally.^17,18^ However, AAV gene therapy has also been associated with some important drawbacks, including the risk of insertional mutagenesis, potentially-fatal liver toxicity, and strong immunogenicity resulting in an inability to redose.^19,20^

Messenger RNA (mRNA)-based drugs are an emerging class of therapeutics that have the potential to restore wildtype protein levels and function in the context of inherited diseases, without the genotoxicity concerns associated with viral DNA delivery.^21–23^ Since mRNA therapy involves supplying cells with a single-stranded mRNA template and relying upon the ribosomal machinery to produce fully functional bioactive proteins *in situ*, it enables the replacement of large, complex proteins such as transmembrane proteins.^24^ Preclinical studies have demonstrated the efficacy of mRNA-based therapeutics across a host of different genetic diseases, including cystic fibrosis,^24^ alpha 1-antitrypsin deficiency,^25^ methylmalonic acidemia,^26^ and hemophilia B,^27^ among others. Clinical trials are ongoing to investigate the efficacy of mRNA for the treatment of propionic acidemia (ClinicalTrials.gov Identifier: NCT04159103) and cystic fibrosis (ClinicalTrials.gov Identifier: NCT03375047).

Structurally, an *in vitro* transcribed (IVT) mRNA molecule comprises multiple components–a 5’ cap, a protein-coding sequence flanked by 5’ and 3’ untranslated regions (UTRs), and a 3’ poly(A) tail.

Each of these components can be modified to enhance the intracellular stability, biocompatibility, and translational capacity of the mRNA.^22^ Systematic efforts have revealed strategies for optimizing the 5’ cap,^28^ the 5’ UTR,^29,30^ the 3’ UTR,^30,31^ and the poly(A) tail length.^32^ For the coding sequence, codon optimization has been shown to improve the translatability and structural stability of mRNA molecules.^33,34^ Furthermore, chemical base modification has been shown to enhance mRNA translation while simultaneously reducing immune activation.^35–38^ However, differing approaches to mRNA codon optimization,^22^ coupled with conflicting reports that suggest chemical base modification may not even be necessary for applications requiring repeat dosing,^34^ have contributed to ongoing debate around which mRNA engineering strategies are most effective. To this end, we sought to investigate the efficacy of different codon optimization and base modification strategies *in vitro* using luciferase reporter mRNAs (Figure 1A). We subsequently applied our findings to the engineering of NPC1 mRNA, with a view to investigating whether exogenous mRNA might be useful for the replacement of NPC1 protein in patient fibroblasts. To the best of our knowledge, this is the first demonstration of engineered mRNA being used to correct NP-C1 disease *in vitro*.

**Figure 1.**
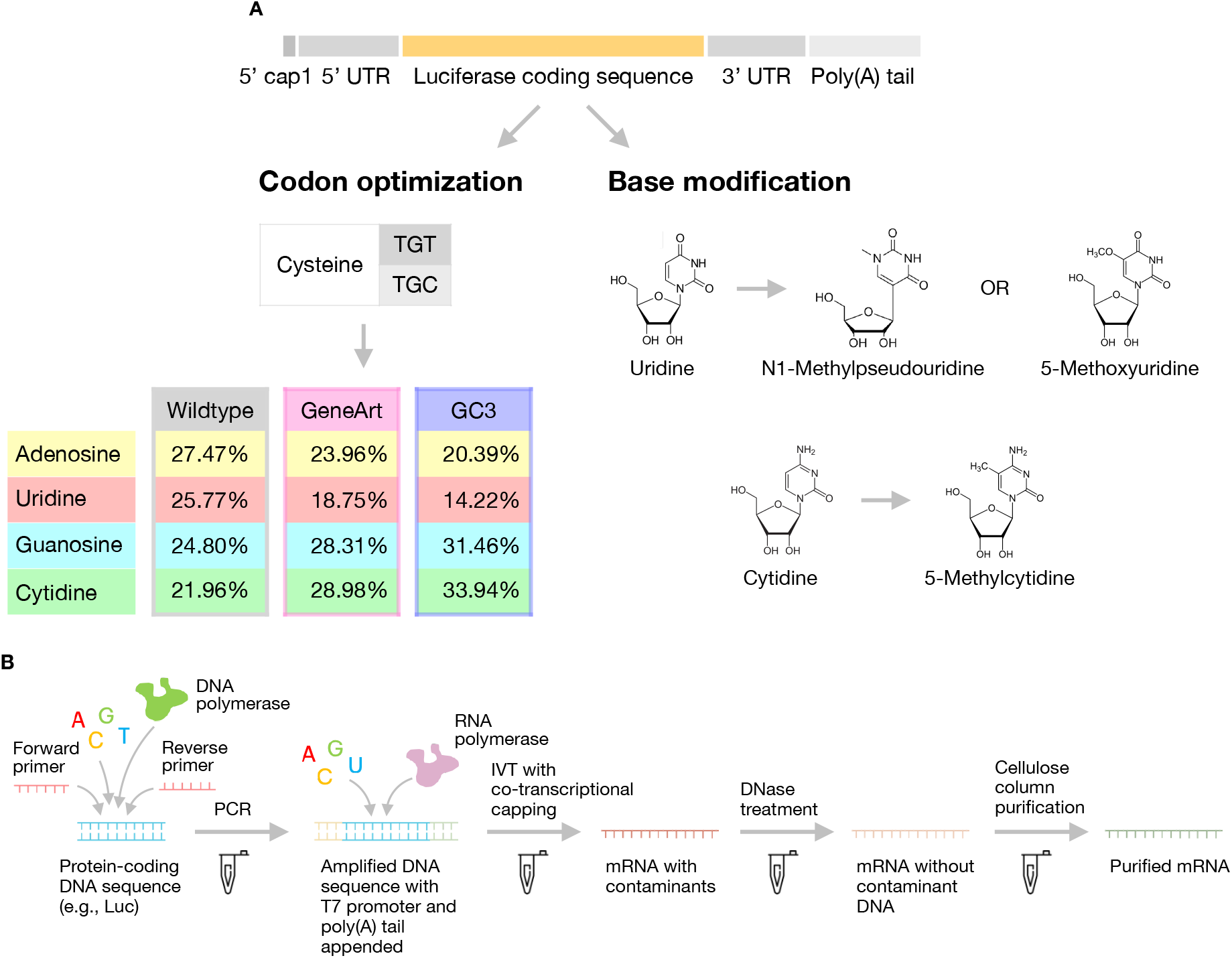
Overview of mRNA engineering workflow. (A) Codon optimization and chemical base modification are two separate engineering strategies that can be used to enhance mRNA translation. Different codon optimization strategies alter the base content of an mRNA coding sequence in different ways. GC3 optimization results in a significant overall enrichment in G and C base content. (B) An overview of the mRNA synthesis and purification workflow used throughout this study.

## Materials and Methods

### Cell culture

HeLa, U87 MG and Hep G2 cells (ATCC) were cultured in Dulbecco’s Modified Eagle’s Medium (DMEM; Lonza) supplemented with 10% fetal bovine serum (FBS; Bovogen), 100 U/ml penicillin and 100 μg/ml streptomycin (Thermo Fisher Scientific). Healthy human control fibroblasts (GM03652G) and NP-C1 patient fibroblasts (GM18393 and GM17919A) were purchased from the Coriell Cell Repository (Table S1) and cultured in DMEM (Lonza) supplemented with 20% FBS (Bovogen), 100 U/ml penicillin and 100 μg/ml streptomycin (Thermo Fisher Scientific). For cholesterol esterification assays, fibroblasts were switched to DMEM supplemented with 5% fetal bovine lipoprotein deficient serum (Alpha Diagnostic International), 100 U/ml penicillin and 100 μg/ml streptomycin. All cell lines were incubated at 37°C with 5% CO_2_.

### mRNA sequence engineering

Codon optimization was implemented to enhance the translatability and structural stability of synthesized mRNAs. The GeneArt-optimized luciferase coding sequence was obtained by inputting the wildtype sequence into the online GeneOptimizer portal (Thermo Fisher Scientific). The GC3-optimized luciferase coding sequence was obtained by substituting, wherever possible, wildtype codons with codons that had a G or C at the third base position. All NPC1 mRNA variants were also codon optimized in the same way (GC3 optimization). This strategy is based on evidence that such codon bias confers enhanced stability and translatability to mRNAs in humans.^39^ Detailed codon selection parameters are available in Supplementary Information (Table S2). All mRNAs harbored an EEF1B2-001 5’ UTR upstream of the open reading frame, and an alpha-1 globin 3’ UTR and 100-nt poly(A) tail downstream of the open reading frame. These sequences have been shown to further enhance the stability and translatability of IVT mRNAs across multiple cell types.^30,40^

### Generation of DNA templates for *in vitro* transcription

Wildtype, GeneArt and GC3 luciferase variants were ordered as separate GenParts DNA fragments (Genscript). The GC3-optimized NPC1 sequence was encoded in a plasmid (Twist Bioscience) and linearized via double restriction enzyme digestion using BamHI-HF (NEB) and HindIII-HF (NEB). Successful linearization was confirmed via 1% agarose gel electrophoresis. Templates for IVT were generated by performing high-fidelity PCR using Q5 Hot Start High-Fidelity DNA Polymerase (NEB) and primers 1 and 2 (IDT; sequences provided in Table S3). From 5’ to 3’, each double-stranded DNA template contained a T7 promoter, a 5’ UTR, open reading frame, 3’ UTR, and 100-nt poly(A) sequence (introduced via primer 2). After linearization and PCR amplification steps, DNA products were purified using the DNA Clean & Concentrator-5 Kit (Zymo Research). All purified PCR products were visually checked for expected size via 1% agarose gel electrophoresis and assessed for purity and yield via Nanodrop (Thermo Fisher Scientific).

### mRNA synthesis and purification

mRNAs were transcribed from DNA templates using T7 RNA polymerase (HiScribe T7 High Yield RNA Synthesis Kit, NEB). To generate base modified mRNA variants, where specified uridine 5’ triphosphate (UTP) was completely replaced with either N1-methylpseudouridine 5’-triphosphate (M1ψTP, APExBIO) or 5-methoxyuridine 5’-triphosphate (5moUTP, APExBIO), and cytidine 5’-triphosphate (CTP) was completely replaced with 5-methylcytidine 5’-triphosphate (5mCTP, APExBIO). Co-transcriptional capping was performed using the EZ Cap Reagent AG (3’ OMe) (APExBIO), resulting in a stable Cap1 structure at the 5’ end of synthesized mRNA transcripts. After IVT reactions had run to completion, DNA templates were digested using DNase I (NEB). mRNAs were subsequently purified using a protocol that has been described previously (Figure 1B).^41^ Purified mRNA was suspended in 1 mM sodium citrate, pH 6.5 (THE RNA Storage Solution, Thermo Fisher Scientific). mRNA integrity was assessed via gel electrophoresis by running denatured aliquots on a non-denaturing 1% agarose gel, using SYBR Safe DNA Gel Stain (Thermo Fisher Scientific) to visualize bands. mRNA concentration and purity were assessed via Nanodrop (Thermo Fisher Scientific).

### Luciferase assay

Cells were seeded at varying densities in white, flat bottom 96-well plates (Corning). The next day, mRNA was complexed with Lipofectamine MessengerMax (Thermo Fisher Scientific) in Opti-MEM I Reduced Serum Medium (Thermo Fisher Scientific) according to the manufacturer’s instructions, using 2 μl Lipofectamine reagent per μg of mRNA. Complexed mRNA was added to cells in serum-containing culture medium. At various timepoints post-transfection (6, 24 or 48 h), plates were retrieved from the incubator and allowed to equilibrate to room temperature, before reconstituted luciferase reagent (ONE-Glo EX, Promega) was added to cells in culture medium, following the manufacturer’s protocol. Plates were incubated at room temperature for 2–3 min with moderate shaking to facilitate cell lysis and sample equilibration. Luminescence was subsequently measured using an Infinite M200 Pro plate reader (Tecan).

### In silico prediction of mRNA secondary structures

mRNA secondary structures were predicted by inputting mRNA sequences into the *RNAfold* webserver provided by the Vienna RNA Websuite.^42^ RNAfold computes the minimum free energy (MFE) and optimal secondary structure for a given sequence using experimentally determined thermodynamic parameters. Default RNA parameters from the 2004 Turner model were used to generate secondary structure predictions.^43^

### Determination of mRNA melting curves via differential scanning fluorimetry

mRNA thermal stability was assessed via differential scanning fluorimetry according to a previously published protocol.^44^ Briefly, 1 μg aliquots of mRNA were mixed with Quant-iT RiboGreen RNA reagent (Thermo Fisher Scientific) and brought to a final volume of 50 μl with nuclease-free water, such that the final RiboGreen concentration was 1X. Samples were added to separate wells of a 48-well PCR plate (Bio-Rad) and incubated in a MiniOpticon Real-Time PCR System (Bio-Rad). A melting curve routine was initiated, whereby samples were incubated at 20°C for 3 min, followed by heating at a constant rate from 20 to 90°C. Fluorescence intensities were recorded at each incremental temperature using the excitation and emission settings for SYBR Green (since RiboGreen and SYBR Green have very similar spectral properties). Once the temperature reached 90°C, samples were cooled to 20°C and allowed to refold for 15 min, at which point the melting curve routine was repeated. Heating and cooling cycles were performed 3 times consecutively, with all samples measured in duplicate. The negative first derivative of fluorescence intensities were then analyzed as a function of temperature.

### NPC1 immunofluorescence staining

Cells were seeded at a density of 3000 cells/well in black, clear-bottom CellCarrier-96 Ultra microplates (PerkinElmer). The next day, cells were transfected with NPC1 mRNA using Lipofectamine MessengerMax reagent, as described above. 24 h after transfection, cells were fixed in 4% paraformaldehyde for 20 min at room temperature, washed with PBS, and permeabilized for 1 h with 0.2% saponin (Sigma-Aldrich) diluted in PBS containing 10% normal goat serum (Thermo Fisher Scientific). Cells were then incubated for 1 h at room temperature with anti-NPC1 primary antibody (ab134113, Abcam) diluted 1:200 in PBS containing 0.2% saponin and 10% normal goat serum. Cells were subsequently washed with PBS containing 0.1% Tween 80, and incubated for 1 h with Alexa Fluor 488-labelled goat anti-rabbit secondary antibody (Thermo Fisher Scientific) diluted 1:500 in PBS containing 0.2% saponin and 10% normal goat serum. After washing with 0.1% Tween 80, cells were stained 1:10000 with Hoechst 33342 (Thermo Fisher Scientific) for 10 min, washed once with PBS, and stored in PBS at 4°C until required for imaging. Cells were imaged using a PerkinElmer Operetta CLS High Content Imaging System with a 20X high numerical aperture dry objective lens and AF488 and DAPI filter sets. At least 7 imaging fields were analyzed per well, and every experimental condition was run in triplicate, equating to an analysis of >300 cells per treatment condition. Images were analyzed using a custom analysis sequence in the PerkinElmer Harmony High-Content Imaging and Analysis Software. Briefly, nuclei were segmented and counted based on Hoechst staining intensities, and intracellular NPC1 staining intensities in each well were separately quantified using the AF488 channel; the resulting per well AF488 intensity values were divided by the number of nuclei in each well to yield the average NPC1 fluorescence per well.

### Amplex Red unesterified cholesterol assay

Cells were seeded at a density of 3000–5000 cells/well in CellCarrier-96 Ultra microplates in DMEM containing 20% FBS. After 24 h, cells were transfected with NPC1 mRNA using Lipofectamine MessengerMax reagent, as described above. 24 or 48 h after transfection, the cell culture medium was removed and cells were washed in PBS twice, before 100 μl reconstituted Amplex Red cholesterol assay reagent (Thermo Fisher Scientific), excluding cholesterol esterase, was added to each well as per the manufacturer’s instructions. After incubation at 37°C for 1 h, the fluorescence intensity in each well was measured (Ex = 560 nm; Em = 590 nm) using an Infinite M200 Pro plate reader.

### alamarBlue cell viability assay

mRNA transfections were carried out in 96-well plates, as described above. 24 or 48 h after transfection, alamarBlue cell viability reagent (Thermo Fisher Scientific) was added to the cell culture medium in a 1:10 ratio (alamarBlue: DMEM), as per the manufacturer’s instructions. Cells were incubated at 37°C in the dark for between 1–4 h, and then the fluorescence intensity in each well was measured (Ex = 560 nm; Em = 590 nm) using an Infinite M200 Pro plate reader.

### Cholesterol esterification assay

Cells were seeded in CellCarrier-96 Ultra microplates at a density of 3000–5000 cells/well in DMEM containing 20% FBS. The next day, cells were switched to DMEM containing 5% fetal bovine lipoprotein deficient serum. After 24 h, cells were transfected with NPC1 mRNA using Lipofectamine MessengerMax reagent, as described above. 24 h after transfection, the culture medium was replaced with either Opti-MEM or Opti-MEM spiked with 50 μg/ml LDL. After a further 6 h, intracellular levels of unesterified cholesterol and total cholesterol were separately assayed using the Amplex Red Cholesterol Assay Kit (Promega), following the manufacturer’s instructions. Levels of esterified cholesterol were calculated by subtracting the amount of esterified cholesterol from the total cholesterol amount. The addition of LDL after 48 h of sterol depletion provides a source of cholesterol that can be used by cells for esterification. While healthy cells are expected to demonstrate a burst of cholesterol esterification in response to this, NP-C patient cells tend to show low levels of esterification regardless of how much cholesterol is available.

### Filipin staining

The same protocol was followed as in the cholesterol esterification assay above. After the 6 h incubation with 50 μg/ml LDL or Opti-MEM alone, cells were fixed in 4% paraformaldehyde for 20 min at room temperature and then washed with PBS. Cells were subsequently stained with 50 μg/ml filipin (Sigma-Aldrich; freshly dissolved in DMSO at 10 mg/ml and then diluted in PBS) in the dark at room temperature for 1 h. After washing, cells were stored in PBS at 4°C, and later imaged using a PerkinElmer Operetta with a 20X high numerical aperture dry objective lens and the DAPI filter set. At least 9 imaging fields were analyzed per well, and every experimental condition was run in triplicate, equating to an analysis of >600 cells per treatment condition. Images were analyzed using a custom analysis sequence in the PerkinElmer Harmony High-Content Imaging and Analysis Software. Intracellular filipin staining intensities in each well were quantified using the DAPI channel.

### Lysotracker Red staining

The same protocol was followed as in the cholesterol esterification assay above. After the 6 h incubation with 50 μg/ml LDL or Opti-MEM alone, cells were live-stained with 50 nM Lysotracker Red DND-99 (Thermo Fisher Scientific) in Opti-MEM at 37°C for 1 h. Cells were subsequently washed with PBS, fixed in 4% paraformaldehyde, and stained with 1:10,000 Hoechst 33342, before being stored in PBS at 4°C. Imaging was performed using a PerkinElmer Operetta with a 40X high numerical aperture dry objective lens and TRITC and DAPI filter sets. At least 9 imaging fields were analyzed per well, and every experimental condition was run in duplicate, equating to an analysis of >300 cells per treatment condition. To measure lysosome size, images were analyzed using a custom analysis sequence in the PerkinElmer Harmony High-Content Imaging and Analysis Software. Briefly, Lysotracker-stained cellular regions were identified and segmented, and the areas of the selected regions were measured and summed for each well.

### Statistics

Experimental data were analyzed used Prism 9 Software (GraphPad), by either ordinary one-way ANOVA with a Dunnett’s multiple comparisons test, or two-way ANOVA with a Tukey’s multiple comparisons test.

## Results

### “GC3” codon optimization enhances mRNA translation, in part due to increased mRNA structural stability

An mRNA molecule’s competence for protein translation is governed by two factors: functional halflife and translational efficiency.^33^ The former is defined by the length of time within which an mRNA molecule is capable of generating protein, while the latter is defined by the mRNA’s ability to recruit ribosomes to initiate translation. The coding sequence plays an important role in shaping both of these factors, and thus governs an mRNA’s translatability to a considerable extent. Codon optimization, the process of replacing wildtype codons with more “optimal” synonymous codons, is a well-known and widely used strategy for enhancing the expression of protein-coding genes.^45^ However, specific strategies for codon optimization vary considerably based on different underlying assumptions about which codon features are important for translation. One approach involves substituting in the most frequently used codon for all instances of a given amino acid.^46^ Another approach involves only replacing rare codons with more abundant synonymous codons.^47^ Still other approaches involve adjusting the codon usage frequency to match the natural frequencies in a host organism,^48,49^ or choosing codons based on cognate transfer RNA (tRNA) abundance.^50^ Many available codon optimization algorithms involve a weighted combination of two or more different optimization parameters.^46,49,51^

A recent study by Hia *et al*. revealed that human mRNAs comprising codons with G or C at the third base position (GC3) are associated with increased stability relative to mRNAs harboring codons with either A or U at the third base position.^39^ They proposed that GC3 codons contribute to mRNA stability in part by preventing interactions with RNA-binding proteins that typically act to induce transcript decay (such as ILF2 and ILF3).^39^ In light of this finding, as well as other corroborating reports,^34,52^ we sought to explore whether GC3 optimization is a viable strategy for enhancing heterologous mRNA expression *in vitro*. To this end, we engineered a firefly luciferase (Luc) coding sequence based on the GC3 optimization strategy, globally optimizing every amino acid in the sequence. Where 2 or more synonymous GC3 codons existed, we made our selection on the basis of which codon had the highest human codon usage frequency. We sought to compare the expression profile of GC3-optimized Luc to wildtype Luc, as well as a Luc variant optimized by a different codon optimization strategy (for the purpose of this study, we used the GeneOptimizer algorithm provided by Thermo Fisher via its GeneArt gene synthesis service).^51^ The GeneOptimizer algorithm takes a multiparameter sliding-window approach to codon optimization, balancing various considerations including GC content, DNA motifs, and codon usage. We designed all 3 mRNAs to harbor a Cap1 structure and EEF1B2-001 5’ UTR immediately upstream of the coding sequence, and an alpha-1 globin 3’ UTR and 100-nt poly(A) tail downstream. Both the EEF1B2-001 5’ UTR and alpha-1 globin 3’ UTR were chosen based on previous reports that showed their inclusion significantly enhanced mRNA stability and expression *in vitro*.^30,40^

We initially produced the 3 Luc mRNAs using all unmodified nucleotides, following a uniform synthesis and purification protocol (Figure 1B). After transfecting 3 different cell lines (we used 3 different cell lines to ensure that our findings were not cell line-specific), we observed that both codon-optimized Luc variants consistently outperformed wildtype Luc by more than an order of magnitude (Figure 2A). However, we observed no meaningful difference in expression profiles between the 2 codon optimization strategies. Given that base modification has been shown to enhance mRNA translation while reducing immune activation,^35,53^ we next investigated the expression profiles of the 3 Luc variants when uridine triphosphate (UTP) was globally replaced with N1-methylpseudouridine triphosphate (M1ψTP). Again, we observed that both codon-optimized variants outperformed wildtype Luc across the 3 different cell lines, while all three variants outperformed their unmodified counterparts (Figure 2B). This time, however, we observed that GC3-optimized Luc produced meaningfully greater protein expression than GeneArt-optimized mRNA.

**Figure 2.**
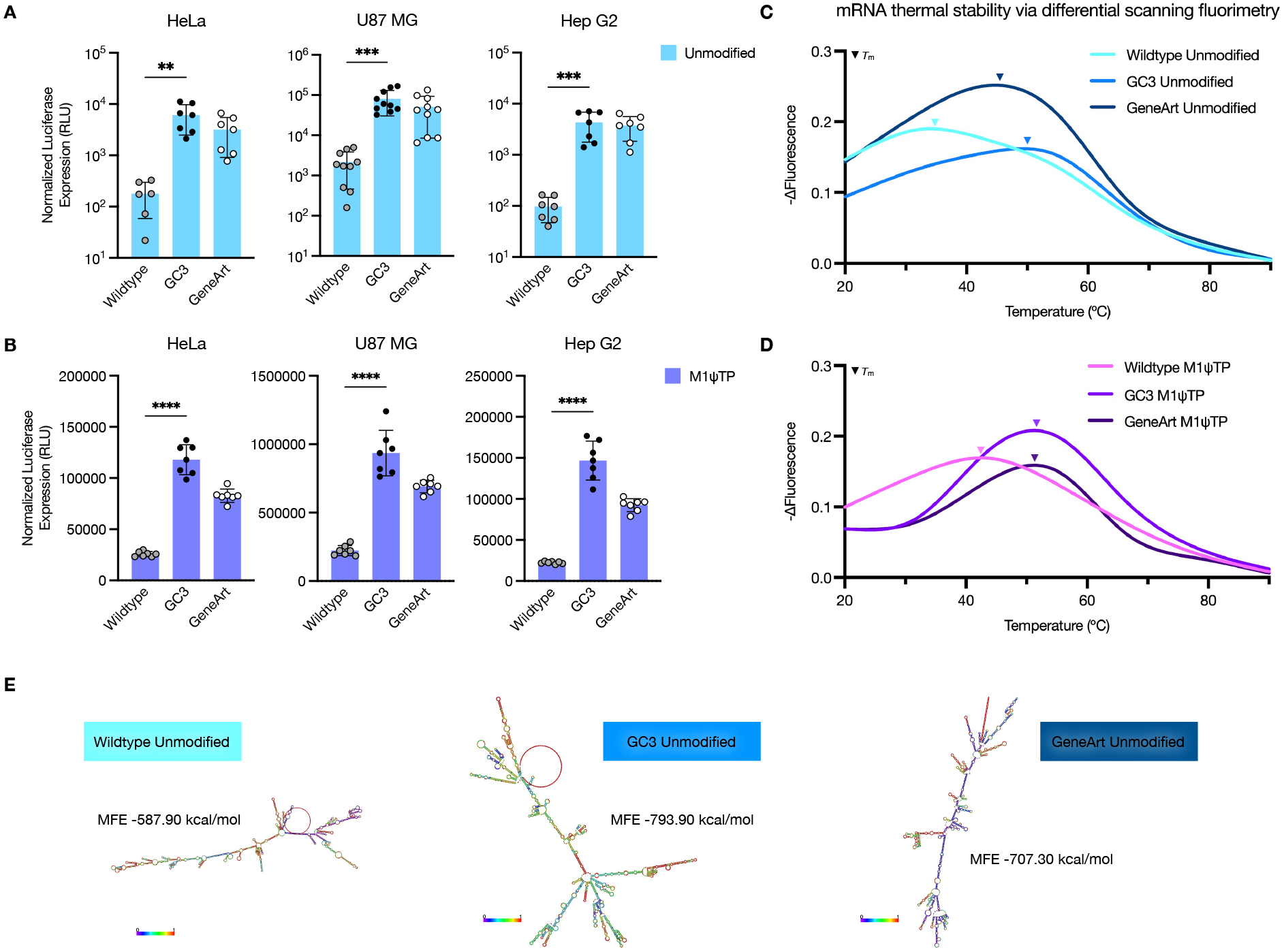
*In vitro* expression and thermal stability profiles of wildtype versus codon-optimized Luc mRNAs. (A) Luciferase expression profiles for Luc mRNA variants harboring all unmodified bases, across HeLa, U87 MG, and Hep G2 cell lines. (B) Protein expression profiles for N1-methylpseudouridine triphosphate (M1ψTP) base-modified Luc mRNA variants, across HeLa, U87 MG, and Hep G2 cell lines. (C, D) Thermal stability curves for unmodified and M1ψTP-modified mRNA variants, as measured by differential scanning fluorimetry. Curves were generated by plotting the negative first derivative of the fluorescence trace as a function of temperature. mRNA melting temperature is denoted by the ▼ icon. (E) RNAfold predictions of optimal secondary structure and minimum free energy for unmodified mRNA variants. Colors indicate base-pairing probabilities, with blue indicating lower probability. In (A) and (B), cells were seeded at a density of 10,000 per well, transfected with 200 ng mRNA, and assayed for protein expression 24 h after treatment. Data are mean ± s.d. of at least three independent experiments and were analyzed by ordinary one-way ANOVA followed by a Dunnett’s multiple comparisons test. \*\**P* < 0.01, \*\*\**P* < 0.001, and \*\*\*\**P* < 0.0001. In (C), all mRNAs were measured in duplicate and for three consecutive heating/cooling cycles. Three independent runs were performed. Curves represent the mean of all independent runs, using data taken from the second heating/cooling cycle.

Having confirmed that GC3 optimization is a viable strategy for enhancing mRNA translation *in vitro*,we next sought to investigate why this might be the case. A previous report by Mauger *et al*. showed that increased mRNA secondary structure leads to enhanced protein expression via an increase in mRNA functional half-life.^33^ Since an mRNA’s GC content tends to be positively correlated with its degree of secondary structure, we probed whether GC3 optimization enhances the secondary structure of Luc mRNA. We used differential scanning fluorimetry, a validated technique that uses an RNA-specific fluorescent reporter dye to measure the structural stability of RNA constructs in the presence of altered environmental parameters such as temperature, salts or pH.^44^ We generated melt curves for the different mRNA variants, measuring each variant during three consecutive cycles of gradual heating from 20°C to 90°C followed by gradual cooling back to 20°C. Using data from the second heating/cooling cycle (due to the greater consistency of the data across several experimental runs compared to the first and third cycles), we determined the peak melting temperature (*T_m_*) for each variant. Consistent with the protein expression data, we observed that both GeneArt unmodified Luc and GC3 unmodified Luc underwent melting at considerably higher temperatures (~46°C and ~51°C) than unmodified wildtype Luc (~35°C), indicating greater mRNA structural stability (Figure 2C). This trend also held for M1ψTP modified constructs, although base modification appeared to dramatically increase the *T*_m_ of wildtype Luc (~35°C to ~43°C) while inducing a much less significant increase for codon-optimized mRNAs (Figure 2D). An estimation of the minimum free energy (MFE) and optimal secondary structure of the unmodified Luc variants using the *RNAfold* webserver^42^ further corroborated these findings (Figure 2E), with wildtype Luc estimated to have a considerably higher MFE than GeneArt and GC3 Luc (a lower MFE indicates greater structural stability). Together, these data suggest that codon optimization–and in particular GC3 optimization–results in increased mRNA stability due to an increase in secondary structure, and that this may play an important role in enhancing protein expression. The possible mechanism(s) by which mRNA secondary structure enhances translation have been discussed elsewhere.^33^

### GC3 codon optimization and N1-methylpseudouridine base substitution synergistically enhance mRNA translation

Next, we investigated the impact of different mRNA base modification strategies on protein expression *in vitro*. M1ψTP and 5-methoxyuridine triphosphate (5moUTP) are among the most commonly used UTP substitutes, while 5-methylcytidine triphosphate (5mCTP) is sometimes used in place of cytidine triphosphate (CTP) and in combination with M1ψTP.^53,54^ Thus, we engineered GC3 and GeneArt Luc mRNAs each with 4 different base compositions: (1) all unmodified bases, (2) 5moUTP in place of UTP, (3) M1ψTP in place of UTP, and (4) M1ψTP in place of UTP and 5mCTP in place of CTP. We transfected HeLa, U87 MG and Hep G2 cells with these 8 mRNA variants and assayed Luc expression at 24 h. All 3 cell lines exhibited relatively uniform expression profiles, with the different base modification strategies ranking, in order of Luc expression from highest to lowest, M1ψTP > 5moUTP > M1ψTP + 5mCTP > unmodified (Figure 3A). We observed that the combination of GC3 codon optimization with M1ψTP base modification cumulatively resulted in the highest protein expression across all cell lines. M1ψTP has been shown to enhance mRNA translation by increasing ribosome recruitment and thus improving translational efficiency.^35^ Time course experiments revealed that kinetic Luc expression profiles varied across the three cell lines, with expression peaking at 6 h in HeLa cells but at 24 h in U87 MG and Hep G2 cells (Figure S1). Interestingly, although GC3 optimization outperformed GeneArt optimization across both 5moUTP and M1ψTP modifications, the same did not hold for the 5mCTP base modification strategy (GeneArt optimization outperformed GC3 optimization). The dramatic reduction in expression when 5mCTP was combined with M1ψTP, relative to M1ψTP alone, has previously been observed in cells and cell-free extracts.^35^ Thus, we conclude that coupling GC3 optimization with M1ψTP base modification leads to synergistic enhancement of mRNA translation *in vitro*. In our hands, this combination generated an mRNA that was approximately 1000-fold more potent than wildtype, unmodified Luc mRNA (Figure 3B).

**Figure 3.**
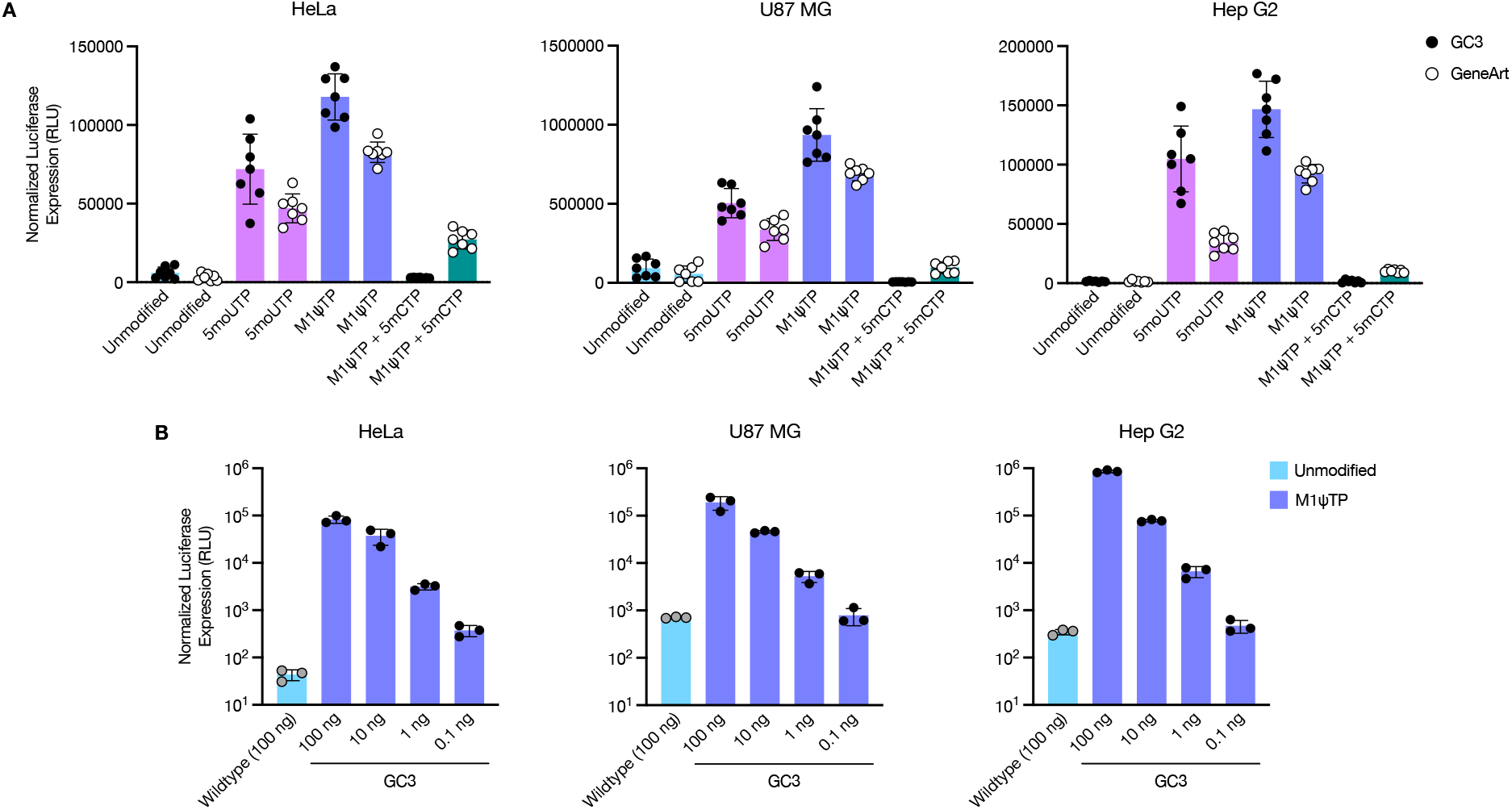
*In vitro* expression profiles for codon-optimized Luc mRNAs harboring different base modifications. (A) Protein expression profiles for GC3-optimized and GeneArt-optimized Luc mRNAs harboring either all unmodified bases, M1ψTP-modified bases, 5moUTP-modified bases, or M1ψTP- and 5mCTP-modified bases. Expression was assayed across HeLa, U87 MG, and Hep G2 cell lines. (B) Protein expression profiles for GC3-optimized, M1ψTP-modified Luc mRNA at 100 ng, 10 ng, 1 ng, and 0.1 ng, compared to wildtype, unmodified Luc mRNA at 100 ng. In (A), cells were seeded at a density of 10,000 per well, transfected with 200 ng mRNA, and assayed for protein expression after 24 h. Data are mean ± s.d. of at least three independent experiments. In (B), HeLa and U87 MG cells were seeded at a density of 10,000 per well, while Hep G2 cells were seeded at a density of 15,000 per well. Protein expression was assayed 24 h post-transfection. Data are mean ± s.d. of three technical replicates, from one representative of three independent experiments.

### Engineered NPC1 mRNA restores functional protein expression and reverses disease pathology in NP-C1 patient fibroblasts

Having successfully engineered a highly expressing Luc mRNA, we next sought to apply our findings to the engineering of NPC1 mRNA (Figure 4A). We performed GC3 optimization of the NPC1 coding sequence and appended the same 5’ and 3’ flanking regions as before to enable efficient protein expression. We synthesized 3 NPC1 mRNA variants, all comprising identical sequences but utilizing different base compositions (unmodified, M1ψTP-modified, and 5moUTP-modified), to test whether the same trend that we observed in reporter assays extended to a human *in vitro* disease model. We did not synthesize constructs with 5mCTP due to the dramatic decrease in overall expression that we observed in Luc reporter assays when 5mCTP base modification was combined with M1ψTP modification. To determine whether the NPC1 mRNA constructs could induce functional protein expression *in vitro*, we transfected them into human fibroblasts derived from two clinically affected NP-C1 patients (GM18393 and GM17919), both which were previously shown to harbor compound heterozygous mutations at the NPC1 gene locus, with different exons affected for each donor. All 3 NPC1 mRNA variants generated significantly increased protein expression in both cell lines compared to the Lipofectamine-treated control, as measured by high-content widefield fluorescence imaging (Figures 4B, S2A and S2B). Having established that all 3 mRNA variants could induce significant NPC1 protein expression *in vitro*, we performed all subsequent experiments using NPC1 M1ψTP mRNA alone due to our previous findings with the Luc mRNA variants. Additionally, due to an insufficiency of GM17919 cells (the growth and viability of these cells became compromised during passaging, despite supplementation with 20% FBS), we performed all further experiments with only GM18393 cells. Automated quantification of NPC1 protein expression via PerkinElmer Harmony High-Content Imaging and Analysis Software revealed that treatment with NPC1 M1ψTP mRNA boosted NPC1 protein levels to the same as those observed in healthy, wildtype donor fibroblasts (Figure 4C).

**Figure 4.**
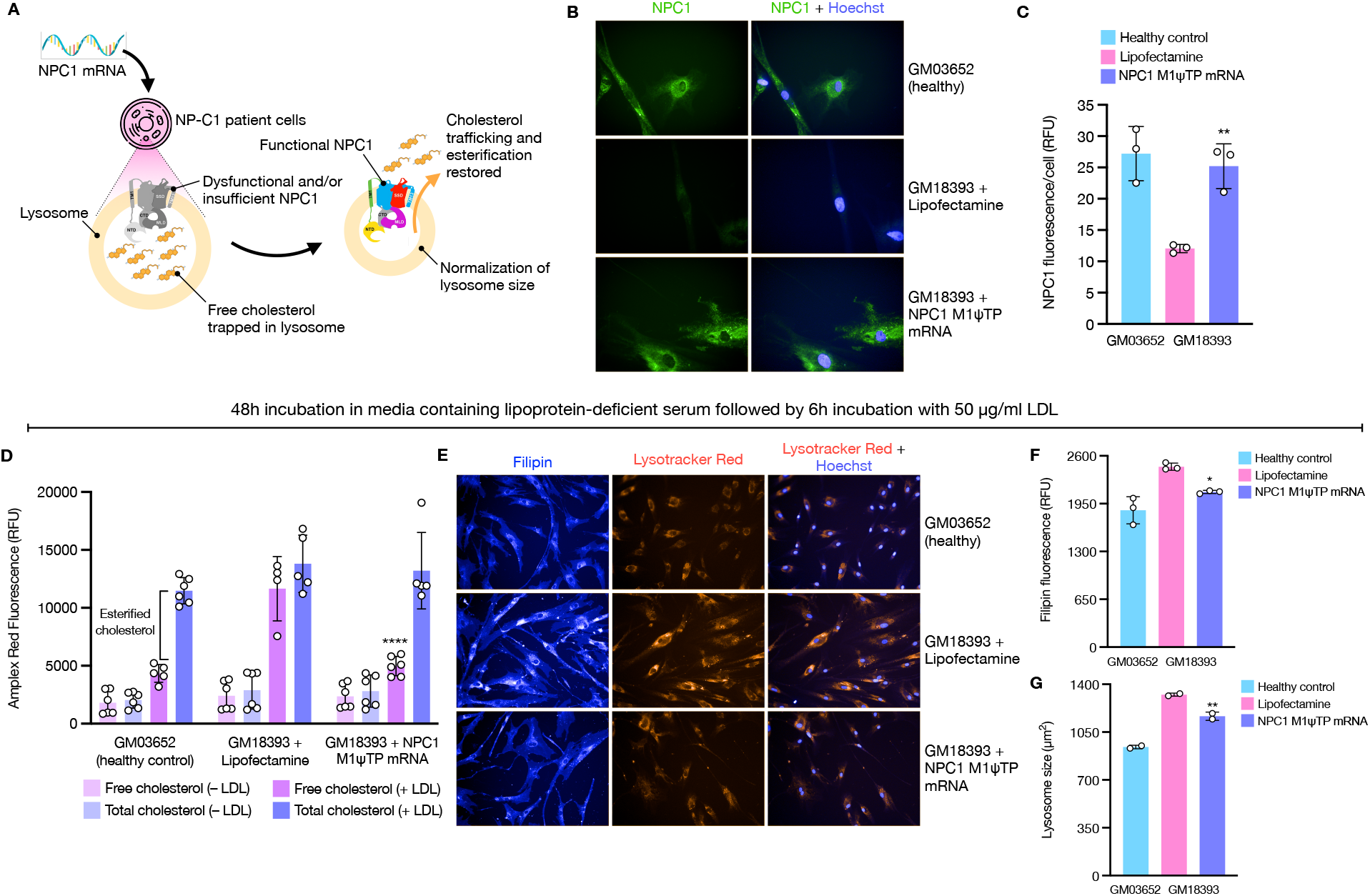
mRNA treatment rescues NPC1 protein insufficiency and the disease phenotype in patient fibroblasts. (A) Overview of the rationale behind mRNA-based treatment of NP-C1 disease. (B) NPC1 and Hoechst 33342 staining in healthy fibroblasts (GM03652), Lipofectamine-treated patient fibroblasts (GM18393), and GM18393 patient fibroblasts treated with 50 ng of M1ψTP-modified NPC1 mRNA. Cells were seeded at a density of 3000 per well and imaged 24 h after treatment. Images are representative of three independent experiments. (C) Quantification of NPC1 staining from the experiment in (B), as measured by average NPC1 fluorescence per cell (cumulative NPC1 fluorescence was divided by the total number of cells identified via Hoescht staining, using the PerkinElmer Harmony high-content analysis software). Data are mean ± s.d. of three technical replicates (from one representative of three independent experiments), encompassing 7 imaging fields per well and >300 cells per treatment condition. (D) Quantification of total and unesterified cholesterol levels in healthy cells, Lipofectamine-treated GM18393 cells, and GM18393 cells treated with 50 ng NPC1 M1ψTP mRNA, after a 48 h incubation in lipoprotein-deficient serum followed by either a 6 h incubation with 50 μg/ml LDL or a further 6 h incubation with lipoprotein-deficient serum to establish a baseline control. Total and unesterified cholesterol levels were assayed using the Amplex Red Cholesterol Assay Kit. Data are mean ± s.d. of two independent experiments. (E) Filipin and Lysotracker Red DND-99 staining in healthy cells, Lipofectamine-treated GM18393 cells, and 50 ng mRNA-treated GM18393 cells after a 48 h incubation in lipoprotein-deficient serum followed by 6 h incubation with 50 μg/ml LDL. Images are representative of three independent experiments. (F) Quantification of filipin staining from the experiment in (E), as measured by average filipin fluorescence per well. Data are mean ± s.d. of three technical replicates (from one representative of three independent experiments), encompassing 9 imaging fields per well and >600 cells per treatment condition. (G) Quantification of lysosome size from Lysotracker Red staining in (E), using the PerkinElmer Harmony high-content analysis software. Data are mean ± s.d. of two technical replicates (from one representative of three independent experiments), encompassing 9 imaging fields per well and >300 cells per treatment condition. In (C), (F) and (G), \**P* < 0.05, \*\**P* < 0.01 by ordinary one-way ANOVA followed by a Dunnett’s multiple comparisons test. In (D), \*\*\*\**P* < 0.0001 by two-way ANOVA followed by a Tukey’s multiple comparisons test.

Next, we evaluated whether the mRNA-induced increase in NPC1 protein levels corresponded to phenotypic rescue. Assaying the levels of free, unesterified cholesterol in treated and untreated cells confirmed that the expressed NPC1 protein was functional, since transfected cells showed a clear, albeit modest, reduction in free cholesterol levels at 48 h post-transfection (Figure S3A). Surprisingly, we observed an inverse dose-response relationship, where a lower mRNA dose (25 ng) corresponded to phenotypic improvement (reduced free cholesterol) whereas a higher mRNA dose (100 ng) produced no such improvement. To further probe the downstream phenotypic consequences of mRNA treatment, we performed a cholesterol esterification assay, which is commonly used to diagnose NP-C.^55,56^ Cells were seeded in full serum media, which was subsequently replaced with media containing lipoprotein-deficient serum the following day; a 48 h incubation in lipoprotein-deficient serum served to deplete cells of cholesterol, after which 50 ug/ml LDL was added directly to the cells and incubated for a further 6 h. With LDL supplementation providing a fresh source of cholesterol available for esterification, a comparison of free and total cholesterol levels at the end of 6 h yielded insight into to how much esterified cholesterol was produced by the cells. Unlike healthy cells, which esterify a considerable proportion of the introduced cholesterol pool, NP-C1 cells are incapable of transporting cholesterol across the lysosomal membrane to be esterified, and thus retain cholesterol only in its free, unesterified form. However, treatment with NPC1 M1ψTP mRNA 20–24 h prior to LDL supplementation restored the capacity for cholesterol esterification in patient cells, as evidenced by a significant reduction in free cholesterol levels (>57% reduction compared to lipofectamine-treated control) without a change in total cholesterol levels (Figure 4D). Widefield fluorescence imaging analysis corroborated these findings. After carrying out the 48 h incubation in lipoprotein-deficient serum followed by 6 h incubation with LDL, we stained cells with either filipin, a dye that binds specifically to free cholesterol,^57^ or Lysotracker Red, a dye that is specific for acidic organelles. While Lipofectamine-treated GM18393 cells showed a stark increase in filipin and Lysotracker Red staining relative to healthy control cells, mRNA treatment caused a clear reduction in staining intensities (Figure 4E). Automated image quantification using the PerkinElmer Harmony High-Content Imaging and Analysis Software confirmed that mRNA treatment caused a statistically significant reduction in filipin intensity, with free cholesterol levels essentially reverting to healthy baseline levels (Figure 4F). We also observed a clear reduction in lysosome size (~157 μm^2^ reduction compared to lipofectamine-treated control), similarly measured by automated quantification of Lysotracker Red staining (Figure 4G). Since NP-C1 disease is marked by lysosomal enlargement due to cholesterol entrapment, these data suggest that mRNA treatment causes functional NPC1 protein to be correctly installed in the endo-lysosomal membrane, resulting in the mobilization of trapped cholesterol and normalization of lysosome size. Taken together, our findings show that mRNA treatment has the capacity to rescue NPC1 protein insufficiency and correct the molecular and phenotypic consequences of biallelic *NPC1* mutations.

## Discussion

The utility of mRNA as a therapeutic strategy for protein replacement has been demonstrated across a wide array of genetic diseases.^24–27,58–64^ Building upon this body of work, we sought to investigate whether mRNA treatment could correct the biochemical and phenotypic hallmarks of NP-C1 disease *in vitro*. We first sought to evaluate codon optimization strategies so that we could consistently engineer mRNAs capable of generating high protein expression across multiple cell types. Utilizing the GC3 codon optimization strategy as informed by previous studies,^39,65^ we consistently observed almost two orders of magnitude greater protein expression in a Luc reporter assay compared to wildtype mRNA when all unmodified nucleotides were used (Figure 2A). Protein expression levels generated by GC3-optimized mRNA were comparable to a Luc mRNA variant codon optimized via a different strategy, suggesting that different codon optimization strategies may be valid as a means for enabling therapeutic protein expression. When we paired codon optimization with base modification, however, GC3 optimization combined with M1ψTP base modification emerged as the clearly optimal strategy for maximizing protein expression (Figure 3A).

The increased expression capacity of the GC3-optimized mRNA variant appears to be, in large part, a result of the enhanced functional half-life of the mRNA molecule,^33^ which in turn is influenced by its secondary structure. Our data show that GC3 codon optimization enhances mRNA thermal stability via an increase in secondary structure (Figures 2C and 2D), while M1ψTP base modification also modulates secondary structure and thus thermal stability to a certain extent. The precise relationship between mRNA secondary structure and protein levels remains to be fully elucidated but appears to be governed by multiple mechanisms. GC3 optimization has previously been shown to increase mRNA functional half-life by preventing or reducing interactions with RNA-binding proteins responsible for mRNA degradation, including ILF2 and ILF3.^39^ Additionally, enhanced mRNA secondary structure may increase association with stabilizing RNA-binding proteins.^66,67^ Ribosomal translocation along the open reading frame (ORF) may be positively affected by increased mRNA secondary structure, with both the speed of ribosome movement and frequency of pausing favoring increased protein output versus mRNAs containing a lesser degree of secondary structure.^33^ Finally, the depletion of uridine throughout the ORF as a result of GC3 optimization may further enhance expression by reducing activation of RNA-dependent pathways that lead to inhibition of translation,^52^ such as the RNase L pathway.^68^ Incorporation of M1ψTP further enhances mRNA translatability independent of GC3 optimization, not only by increasing mRNA stability, but also by preventing protein kinase R-induced translational repression and by increasing ribosomal density.^33,35^ Conversely, our finding that 5mCTP base modification severely compromised protein expression, especially when combined with GC3 optimization (Figure 3A), may suggest an adverse effect of 5mCTP incorporation on mRNA secondary structure and ribosome kinetics.^35^ Together, these observations reveal how GC3 optimization coupled with M1ψTP base modification synergistically enhance protein production in cells. One practical constraint surrounding the GC3 optimization strategy, however, is that current commercial gene synthesis technologies are often incapable of producing genes with overtly high levels of complexity (i.e., high GC content). In view of this, the GC3 optimization strategy sometimes needs to be tailored to ensure that a fair tradeoff is achieved between compliance with technology constraints, and maintenance of a relatively high degree of mRNA secondary structure and codon optimality.

Having identified a viable strategy for engineering highly-expressing and minimally immunogenic^35^ mRNAs, we next investigated whether engineered NPC1 mRNA could rescue NP-C1 disease in patient fibroblasts. We first assayed NPC1 protein expression 24 h after mRNA treatment, and observed that all 3 mRNA variants, which differed only in the type of base modification employed (unmodified, M1ψTP, 5moUTP), were able to generate substantial NPC1 protein expression *in vitro* as determined by widefield fluorescence imaging (Figures 4B, S2A and S2B). Since GC3 M1ψTP Luc mRNA generated the best dose-response profile in Luc reporter assays, we decided to narrow our focus to this engineering strategy in all subsequent NPC1 experiments. Our finding that treatment with NPC1 M1ψTP mRNA restored NPC1 protein levels to those observed in normal, healthy cells (Figure 4C) encouraged us to assess the phenotypic consequences of mRNA treatment. We evaluated whether NPC1 M1ψTP mRNA could reduce the levels of free, unesterified cholesterol in affected cells seeded in full serum medium. Although we observed a statistically significant reduction in free cholesterol levels at 48 h post-treatment (Figure S3A), the change was modest. It is possible that the paucity of phenotypic rescue may have been caused by the rapid growth and division of cells in the time that elapsed after mRNA treatment, especially given the high serum content of the media. It is also possible that the selected timepoint may have been shorter than was necessary to observe a more significant phenotypic change.

More surprising, however, was our finding that only the lowest mRNA dose (25 ng) had any meaningful impact on free cholesterol levels, whereas higher doses failed to elicit any substantial phenotypic change. This inverse dose-response relationship was also observed in other phenotypic assays. Filipin imaging in the context of a cholesterol esterification assay revealed that whereas the lowest mRNA dose (50 ng) yielded a phenotypic improvement (a reduction in filipin levels), the highest mRNA dose (200 ng) produced an increase in filipin staining well above the Lipofectamine-treated control (Figure S3B). (Reasons for differential mRNA dosing between assays include differences in seeding densities, growth medium and assay timepoints.) The worsened phenotype at higher doses may have occurred due to mRNA overdosing. An overabundance of mRNA would be expected to introduce many more transcripts into cells than the ribosomes could reasonably process, resulting in the association between non-ribosome-bound mRNAs and aggregation-prone proteins, and the subsequent formation of stress granules.^69^ Cellular stress has previously been linked to an upregulation in sterol synthesis and cholesterol accumulation.^70,71^ Since cells affected by NP-C1 lack the ability to esterify cholesterol, the increased cholesterol production as a result of cellular stress would be expected to increase the pool of unesterified cholesterol in the lysosome. Further studies would help to elucidate the precise mechanism by which mRNA overdosing leads to an exacerbation of the NP-C1 disease phenotype. It would also be useful to know whether such a phenomenon is generalizable across disparate disease states.

To further probe whether NPC1 M1ψTP mRNA could rescue the disease phenotype, we subjected cells to a cholesterol esterification assay, depleting cells of cholesterol for an extended period of time before reintroducing cholesterol in the form of LDL and monitoring cholesterol fate after 6 h. Quantification of the levels of free and total cholesterol revealed that, at the appropriate dose, NPC1 M1ψTP mRNA could indeed rescue the disease phenotype, as evidenced by a normalization in the levels of esterified cholesterol where minimal capacity for esterification had previously been observed (Figure 4D). Filipin and Lysotracker Red imaging corroborated these findings (Figure 4E), and quantification of filipin levels and lysosome size (the latter of which was measured by tracing the cell regions stained by Lysotracker Red) confirmed a statistically significant reduction of both after mRNA treatment (Figures 4F and 4G). To the best of our knowledge, this is the first time exogenous mRNA has been used to rescue the protein insufficiency and pathogenic phenotype caused by NPC1 disease. Nevertheless, further investigation is required to advance this work towards clinical application. Of significant importance to conducting an *in vivo* proof-of-concept study is the need for a non-viral delivery system capable of safely and effectively transporting NPC1 mRNA to the relevant affected tissues, especially the brain and liver.

## Author contributions

D.F., C.C.-J., F.C., Y.H.H., and A.I.B. designed the research. D.F. conducted the experiments. F.C. supervised the project. D.F. wrote the manuscript with input from all other authors.

## Acknowledgements

This research was funded by the National Health and Medical Research Council (NHMRC) (GNT1149990, F.C.). Y.H.H. and A.I.B. acknowledge grant funding from the Australian NPC Disease Foundation. F.C. acknowledges the award of an NHMRC Senior Principal Research Fellowship (GNT1135806). C.C.-J. acknowledges support from the Friedreich’s Ataxia Research Alliance and fara Australia. D.F. acknowledges the award of a Westpac Future Leader’s Scholarship from the Westpac Foundation. We acknowledge the Biological Optical Microscopy Platform (BOMP) at the University of Melbourne for access to the PerkinElmer Operetta CLS High Content Imaging System.

