## Supplementary Material for "mRNA Treatment Rescues Niemann-Pick Disease Type C1 in Patient Fibroblasts"

**Table S1. Characterization of primary fibroblast cell lines used in this study.**

| Cell line | Identified mutation (allele 1) | Identified mutation (allele 2) |
|---|---|---|
| GM03652 male fibroblast | None (wildtype) | None (wildtype) |
| GM18393 male fibroblast | GLY248VAL (chromosomal location 18q11-q12) | MET1142THR (chromosomal location 18q11-q12) |
| GM17919 female fibroblast | ILE1061THR (chromosomal location 18q11-q12) | ARG404TRP (chromosomal location 18q11-q12) |

**Table S2. Codon selection parameters used for GC3 codon optimization in this study.**

| Amino acid | GC3-optimal codon |
|---|---|
| Alanine (A) | GCC |
| Cysteine (C) | TGC |
| Aspartic acid (D) | GAC |
| Glutamic acid (E) | GAG |
| Phenylalanine (F) | TTC |
| Glycine (G) | GGC |
| Histidine (H) | CAC |
| Isoleucine (I) | ATC |
| Lysine (K) | AAG |
| Leucine (L) | CTG |
| Methionine (M) | ATG |
| Asparagine (N) | AAC |
| Proline (P) | CCC |
| Glutamine (Q) | CAG |
| Arginine (R) | CGG |
| Serine (S) | AGC |
| Threonine (T) | ACC |
| Valine (V) | GTG |
| Tryptophan (W) | TGG |
| Tyrosine (Y) | TAC |

Initial codon optimization was performed by selecting codons with a G or C at the third base position. Where two or more GC3-optimal codons existed, we performed a secondary selection on the basis of which codon had the highest human codon usage frequency. For certain genes, GC3 codon optimization renders the gene too complex for synthesis via commercial gene synthesis technologies. Thus, the GC3 codon selection parameters sometimes need to be relaxed to allow usage of GC3-nonoptimal codons, such that genes can be fully synthesized.

**Table S3. Primers used in this study for generation of DNA templates.**

| Primer # | Sequence (5' to 3') |
|---|---|
| 1 | CCACGAATCTAATACGACTCACTATAAGG |
| 2 | TTTTTTTTTTTTTTTTTTTTTTTTTTTTTTTTTTTTTTTTTTTTTTTTTTTTTTTTTTTTTTTTTTTTTTTTTTTTT TTTTTTTTTTTTTTTTTTTTTTTGCCGCCCACTCAGACTTTATTC |

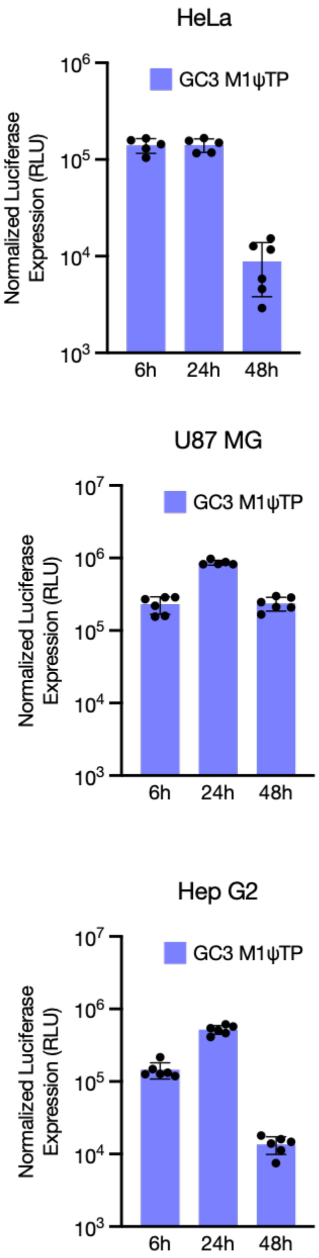


**Figure S1. Luciferase time-course assays for GC3 M1ψTP Luc mRNA across different cell lines**
Luciferase expression at 6 h, 24 h, and 48 h post-transfection with M1ψTP-modified GC3 Luc mRNA in (A) HeLa cells, (B) U87 MG cells, and (C) Hep G2 cells.

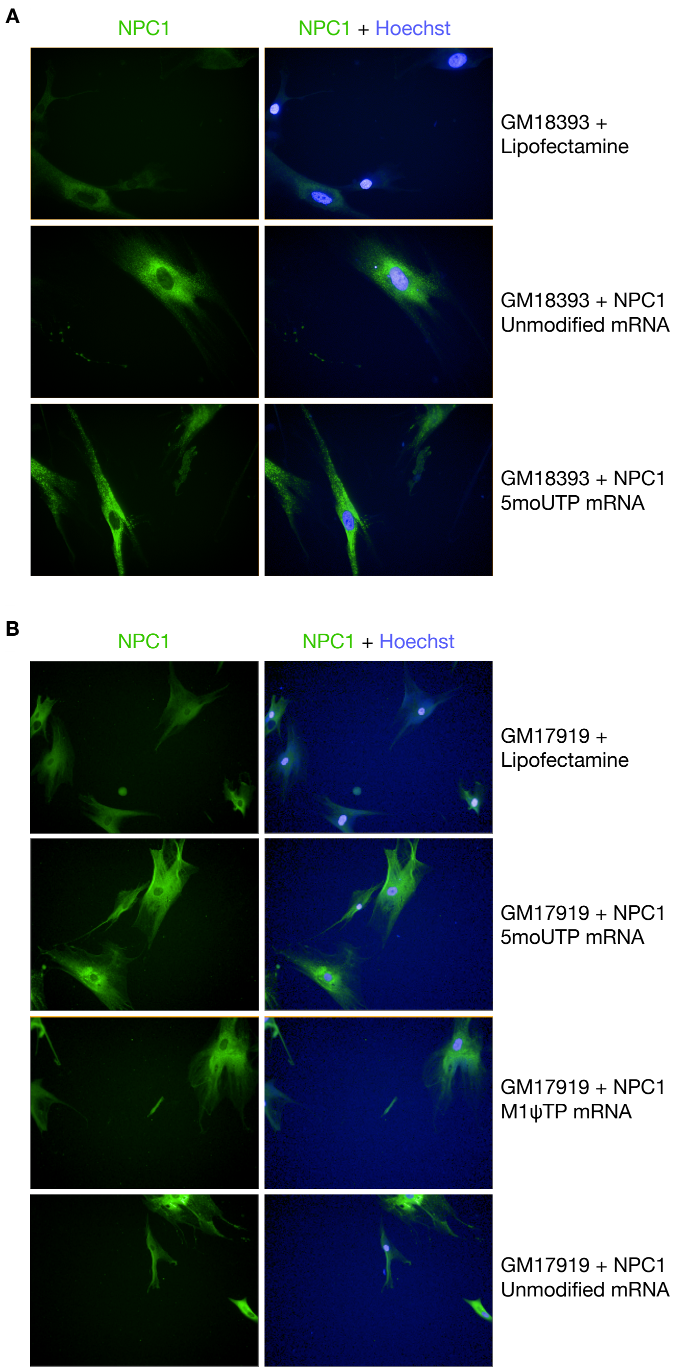


**Figure S2. Transfection with engineered NPC1 mRNA increases NPC1 protein expression in fibroblasts derived from two different NPC1 patients**

(A) NPC1 and Hoechst 33342 staining in GM18393 patient fibroblasts treated with Lipofectamine, 50 ng of unmodified NPC1 mRNA, or 50 ng of 5moUTP-modified NPC1 mRNA. Cells were seeded at a density of 3000 per well and imaged 24 h after treatment. Images are representative of three independent experiments. (B) NPC1 and Hoechst 33342 staining in GM17919 patient fibroblasts treated with Lipofectamine, 50 ng of 5moUTP-modified NPC1 mRNA, 50 ng of M1ψTP-modified NPC1 mRNA, or 50 ng of unmodified NPC1 mRNA. Cells were seeded at a density of 3000 per well and imaged 24 h after treatment. Images are representative of two independent experiments.

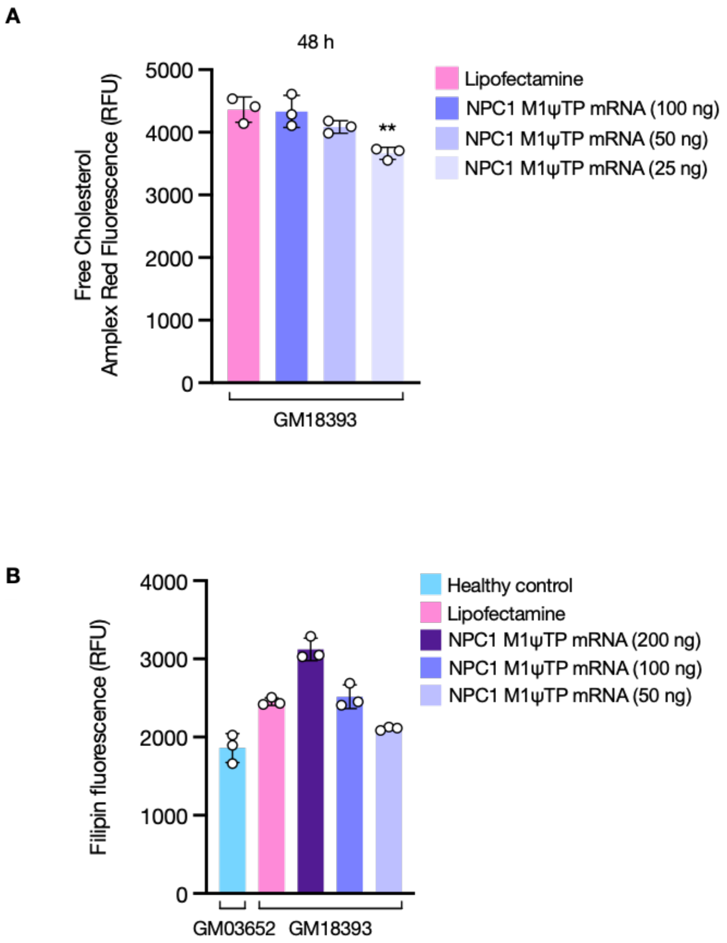


**Figure S3. M1ψTP-modified NPC1 mRNA exerts an inverse phenotypic dose-response effect on GM18393 cells**

(A) Quantification of free, unesterified cholesterol levels in GM18393 cells after treatment with Lipofectamine or three descending doses of M1ψTP-modified NPC1 mRNA (100 ng, 50 ng, 25 ng). Surprisingly, only the 25 ng mRNA dose produced a statistically meaningful reduction in free cholesterol levels. Cells were seeded at a density of 3000 per well in full serum media and assayed 48 h after transfection. Data are mean ± s.d. of three technical replicates, from one of three representative experiments. ***P* < 0.01 by ordinary one-way ANOVA followed by a Dunnett's multiple comparisons test. (B) Quantification of filipin staining in healthy cells (GM03652), Lipofectamine-treated GM18393 cells, and GM18393 cells treated with three descending doses of M1ψTP-modified NPC1 mRNA (200 ng, 100 ng, 50 ng), after a 48 h incubation in lipoprotein-deficient serum followed by 6 h incubation with 50 µg/ml LDL. Data are mean ± s.d. of three technical replicates (from one of three representative experiments), encompassing 9 imaging fields per well and >600 cells per treatment condition. Images were quantified using the PerkinElmer Harmony High-Content Imaging and Analysis Software.
